# Oscillatory sensory stimulation in the delta-band enhances temporal prediction performance

**DOI:** 10.64898/2026.03.29.715181

**Authors:** Peng Wang, Marleen J. Schoenfeld, Alexander Maÿe, Jonathan Daume, Till R. Schneider, Andreas K. Engel

## Abstract

Predicting the time point when an event will occur is fundamental for adaptive behavior, yet it remains unresolved whether temporal prediction can be influenced by low-frequency rhythmic modulation of sensory stimuli. Here, we tested whether external rhythmic sensory stimulation at a frequency in the delta range (0.5 - 3 Hz) alters performance in a visual temporal prediction task. Participants judged whether a moving visual stimulus reappeared too early or too late after disappearing behind an occluder, while the temporal structure of crossmodal sensory input was manipulated across two behavioral sessions. Results indicated that in the visual-auditory conditions, oscillatory stimulation in either the visual or auditory modality improved performance, whereas decaying sensory intensity over time impaired performance. In visual-tactile conditions, oscillatory visual stimulation also enhanced sensitivity, but rhythmic tactile stimulation did not produce a comparable benefit in performance. Critically, tactile stimulation improved performance only when aligned to the expected disappearance of the visual stimulus, demonstrating that the phase relationship between sensory input and intrinsic delta oscillations is behaviorally relevant. Together, these findings indicate that temporal prediction depends on the temporal structure of sensory input and support the relevance of delta-band oscillations in predictive behavior across and within sensory modalities. Hence, rhythmic modulation of sensory stimuli may provide a tool to enhance temporal prediction accuracy by stimulating oscillatory neural dynamics.

## Introduction

Our environment is constantly changing, yet often follows predictable temporal patterns. This allows the brain to anticipate and prepare responses based on temporal regularities^1^. Neural oscillations are pervasive phenomena in the brain^2^, and are thought to play a role in such temporal predictions^3–5^. Previously, oscillations in the alpha band were modified during temporal prediction tasks, when comparing predictive and unpredictive conditions^6,7^. Further, the phase of endogenous oscillations has been found to reset such that neural excitability is elevated in anticipation of an expected stimulus^8^. A recent magnetoencephalography (MEG) study by Daume et al. ^9^ demonstrated these phase adjustments, i.e., enhanced inter-trial phase consistency, during visual temporal prediction tasks in the delta frequency band (0.5 - 3 Hz). These findings were observed in both unimodal visual and crossmodal visual-tactile settings, while no notable power changes in the delta-band were observed, excluding potential confounding effects of evoked responses like contingent negative variation. This raises the question of whether modulating delta oscillations externally could alter temporal prediction performance.

Based on these findings by Daume et al., (2021), we formulated two hypotheses. First, if delta oscillations support temporal prediction, then strengthening them through sensory entrainment should enhance performance. Here, we tested this with oscillatory stimuli designed to entrain endogenous delta oscillations. Second, if phase information is critical, then weakening phase information should impair performance. We tested this with decaying stimuli which conceal stimulus disappearance and thus reduce phase information.

In recent years, modulating neural activity with transcranial alternating current stimulation (tACS) to entrain endogenous oscillations has attracted considerable interest^10–13^. By applying rhythmic external electric fields on the scalp, membrane potentials of neurons are believed to be enhanced at the same frequency^14,15^. However, tACS produces strong and difficult-to-correct electrical artifacts in concurrent EEG and MEG recordings^16,17^. Thus, in this study, we sought to achieve entrainment effects comparable to tACS via sensory inputs, expecting that their induced brain responses would follow the same oscillatory profile and thereby enhance the temporal prediction process. Similarly, a decaying stimulus with ambiguous phase offset was expected to disturb the internal oscillations involved in temporal prediction.

We adopted a temporal prediction task similar to Daume et al. ^9^ in which participants were shown a visual stimulus (an elongated oval) moving horizontally from the periphery toward the center of the screen, disappearing behind an occluder, and reappearing after a certain delay (Figure 1a). Participants were asked to indicate, with a key press, whether the reappearance was “too early” or “too late” relative to their expectation. The experiments were conducted in two sessions (Figure 1b): a visual-auditory (VA) session, in which the movement of the visual stimulus was accompanied by auditory pink noise; and a visual-tactile (VT) session, in which rhythmic tactile stimuli were presented with the moving oval in several conditions. In the VA session, the visual contrast of the moving oval and the volume of the pink noise were modulated to form five conditions: one baseline condition with constant visual contrast and auditory volume (VCAC), and four conditions arranged in a 2×2 design in which each stimulus type was either oscillating or decaying in stimulus intensity: visual oscillating - auditory oscillating (VOAO), visual decaying - auditory decaying (VDAD), visual oscillating - auditory decaying (VOAD), visual decaying – auditory oscillating (VDAO). In the VT session, three visual-only conditions (VCT0, VDT0, VOT0) were presented with either constant (C), decaying (D) or oscillating (O) visual contrast. Two crossmodal conditions (VDT1 and VOT1) included rhythmic tactile stimuli, aligned to the beginning of visual stimuli. In the third crossmodal condition (VDT2), tactile stimuli were phase-shifted, such that the final stimulus was delivered exactly at the disappearance of the oval.

**Figure 1.**
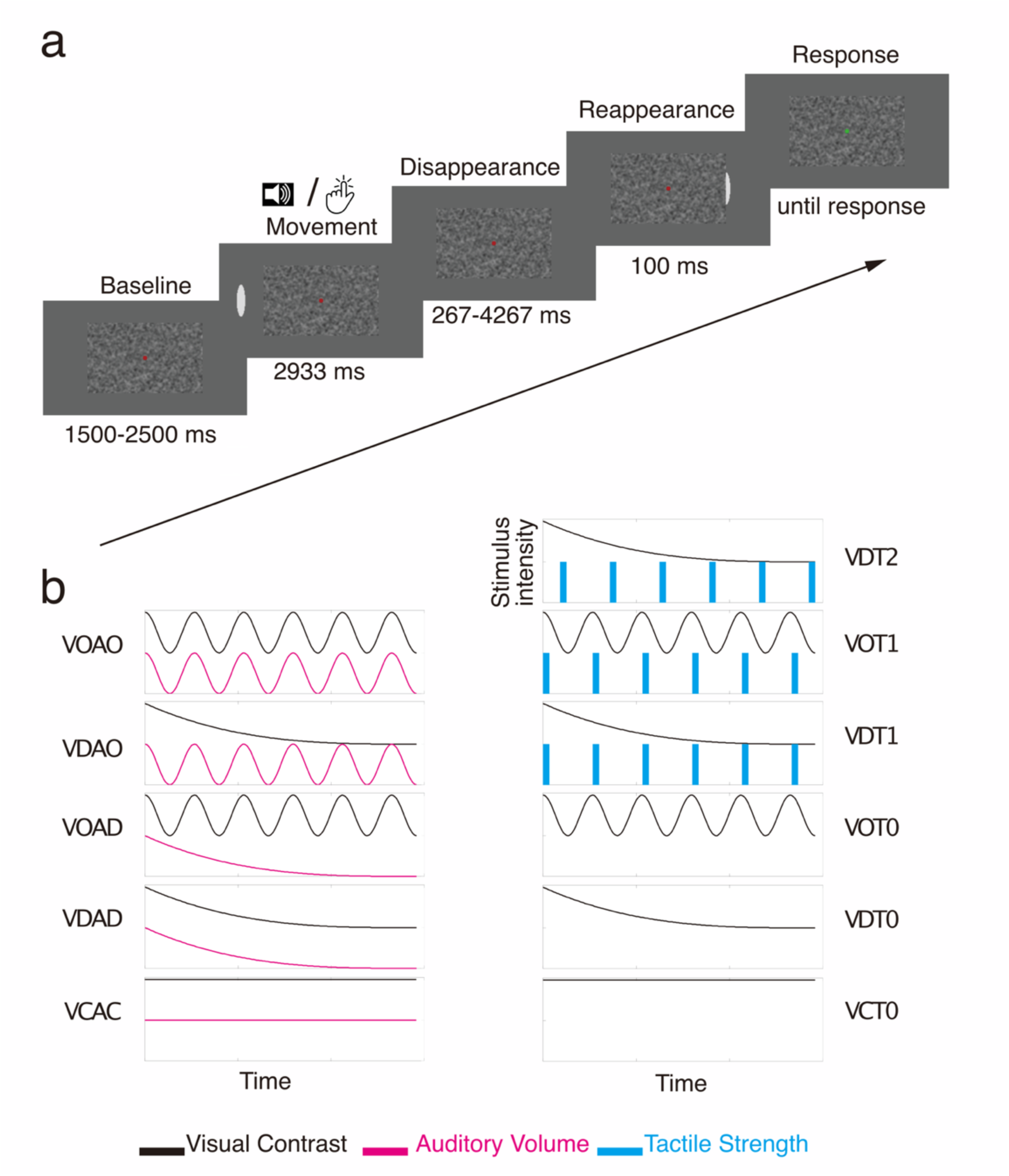
Schematic overview of the experimental design. (**a**) In the task, a red fixation point was displayed at the center of the screen, surrounded by a rectangular area of greyscale white noise (the occluder). A moving visual stimulus (an elongated oval) approached the fixation point, disappeared behind the occluder, and reappeared after a certain delay. Participants were instructed to judge whether the reappearance was too early or too late by pressing a button, when the fixation point turned green. (**b**) The experiment comprised two sessions, one visual-auditory (VA) session (left), in which participants were presented with visual and auditory stimuli and one visual-tactile (VT) session (right), in which participants were presented with visual and tactile stimuli. In the VA session, the moving oval was always accompanied by auditory pink noise, such that the contrast of the oval and the volume of the pink noise were either constant (VC, AC), oscillating (VO, AO) or decaying (VD, AD). This resulted in 5 conditions: One baseline condition, VCAC with constant contrast and volume, and four additional conditions that were designed in a 2 x 2 pattern, in which either stimulus type was oscillating or decaying (VOAD, VDAO) or both were oscillating or decaying (VDAD, VOAO). In the VT session, visual stimuli were again either constant (VC), oscillating (VO) or decaying (VD). Tactile stimuli were either absent (T0), aligned to the onset of the visual stimulus (T1), or aligned to its disappearance (T2).

## Results

Participants were expected to respond “too late” when the reappearance delay exceeded the time required for the visual stimulus to move across the occluder at constant speed, and “too early” otherwise. Through fitting the ratio of “too late” responses across delays with a sigmoid function (equation 1), the parameter ***s***, indicates discrimination sensitivity. Thus, a steeper slope (smaller *s*) indicates more sensitive discrimination between response alternatives, or better performance (see Figure 2a, and methods section for details). The slope ***s*** is used as the primary behavioral metric, similar to previous studies^9^.

**Figure 2.**
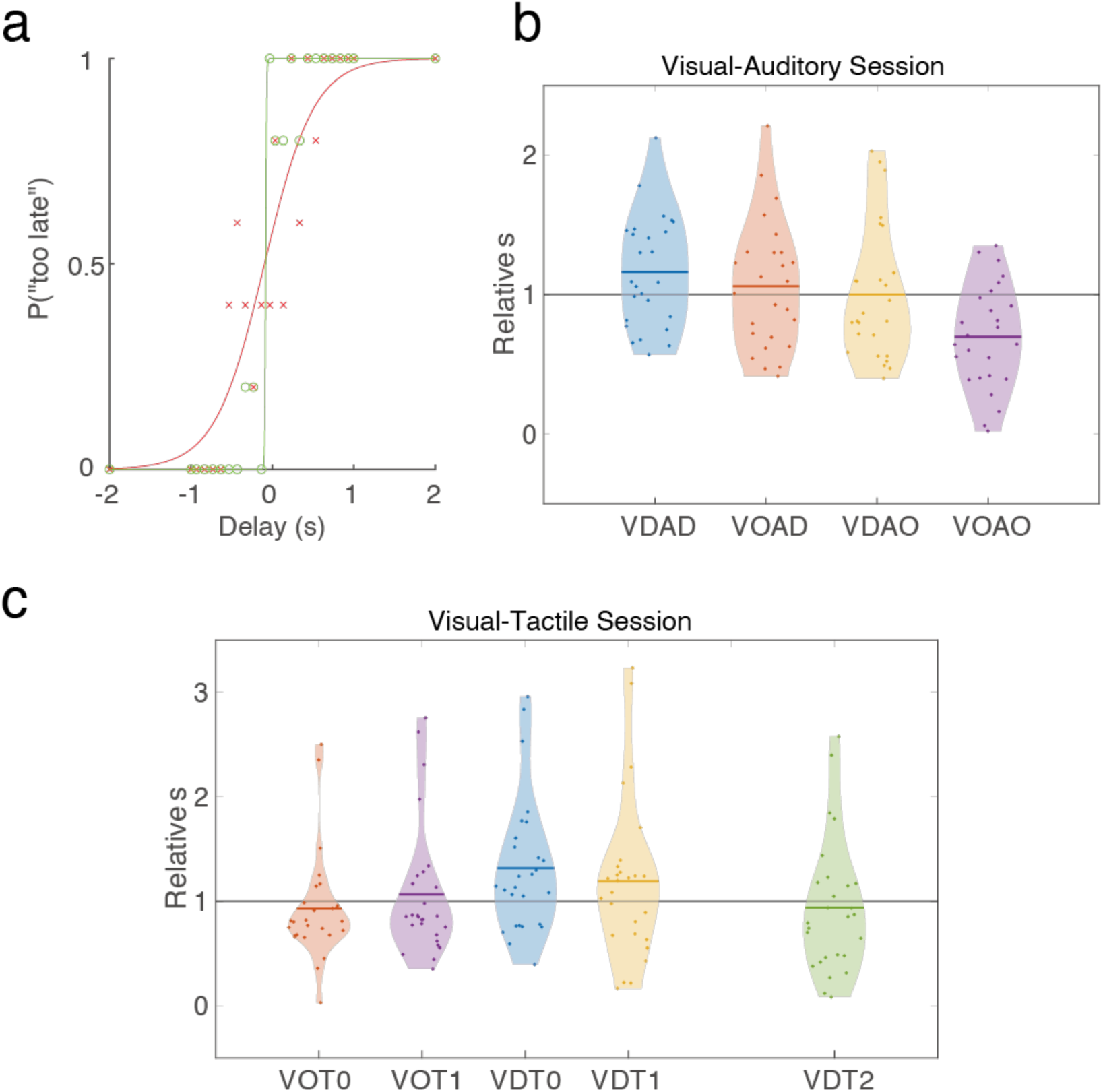
Performance as measured by fitted slope *s*. (c). (**a**) Exemplary results of one participant. The dots indicate the ratio of “too late” responses for the correspondent delays; each represented the average performance of five trials in that delay. They were fitted with a sigmoid function, and the slope *s* was used as a measure of performance: smaller numbers indicate more efficient detection. In this example, the green curve shows a smaller *s* and thus a steeper slope or better performance. (**b**) Results of the visual-auditory session. The slope *s* was normalized to the VCAC condition (horizontal line for relative *s* = 1). (**c**) Results of the visual-tactile session. The slope *s* was normalized to VCT0 condition. VC = visual constant, VD = visual decaying, VO = visual oscillating, AC = auditory constant, AD = auditory decaying, AO = auditory oscillating, T0 = no tactile stimulus, T1 = tactile stimulus not aligned, T2 = tactile stimulus aligned to disappearance of visual stimulus.

### Visual-auditory session

Slope ***s*** obtained from the four conditions VOAO, VDAD, VOAD, and VDAO in the visual-auditory session was normalized by dividing *s* of each condition by *s* of the constant condition (VCAC). To test group differences across conditions, we performed a 2 x 2 ANOVA with two within-subject factors: visual modality and auditory modality, each with two levels: oscillating *vs.* decaying (Figure 2b). No interaction was observed between the visual and auditory factors (*F* (1, 25) = 1.71, *p* = 0.20, *η_p_*^2^ = 0.06). The main effect of the factor visual modality was significant (*F* (1, 25) = 11.72, *p* < 0.01, *η_p_*^2^ = 0.32). Specifically, slopes were steeper in the oscillating condition (mean = 0.88, 95% confidence interval CI = [0.75, 1.01]) compared to the decaying condition (mean = 1.08, CI = [0.94, 1.23]), indicating better performance in the oscillating compared to the decaying conditions. Oscillating conditions also yielded steeper slopes than baseline (*p* = 0.03, *d* = 0.37), indicating better performance compared to baseline. Slopes in the decaying conditions were not significantly different (*p* = 0.11, *d* = 0.26) than baseline. The main effect of the factor auditory modality was also significant (*F* (1, 25) = 15.58, *p* < 0.001, *η_p_*^2^ = 0.39). In particular, the slopes were steeper in the oscillating conditions (mean = 0.85, CI = [0.70, 0.99]) compared to the decaying conditions (mean = 1.11, CI = [0.98, 1.25]), indicating better performance in the oscillating compared to the decaying conditions. Furthermore, the slopes were significantly steeper in the oscillating condition compared to baseline (*p* = 0.02, *d* = 0.42) and shallower in the decaying condition (*p* = 0.05, *d* = 0.33), indicating better and worse performance compared to baseline, respectively.

### Visual-tactile session

The slope ***s*** obtained from the four conditions VDT0, VOT0, VDT1, and VOT1 in the visual-tactile session was normalized by dividing *s* of each condition by *s* of the constant condition (VCT0). To test differences across conditions, we performed a 2 x 2 ANOVA with two within-subject factors visual and tactile modality, each with two levels: oscillating *vs.* decaying for visual stimuli, and on *vs.* off for tactile stimuli (Figure 2c). The results indicated no interaction between visual modality and tactile modality (*F* (1, 26) = 2.74, *p* = 0.11, *η_p_*^2^ = 0.10). In terms of the visual modality, the slope was significantly steeper (*F* (1, 26) = 10.06, *p* < 0.01, *η_p_*^2^ = 0.28) in the oscillating condition (mean = 1.00, CI = [0.78, 1.21]) compared to the decaying condition (mean = 1.25, CI = [1.01, 1.50]), indicating better performance in the oscillating compared to the decaying conditions. The decaying conditions yielded significantly shallower slopes (*p* = 0.02, *d* = 0.42) as compared to baseline, indicating worse performance. No significant difference (*F* (1, 26) = 0.01, *p* = 0.93, *η_p_*^2^ < 0.001) was observed between two levels in the tactile dimension. No difference between the tactile on or off conditions and the baseline was observed either.

Additionally, we tested whether the alignment of the tactile to the visual stimulus influenced performance. To this end, we conducted an ANOVA with one within-subject factor of tactile modality, at three levels: alignment to luminance maximum, to luminance minimum, or absence of tactile stimuli, i.e., VDT1, VDT2, and VDT0 (Figure 2c). The main effect of tactile modality was significant (*F* (2, 25) = 5.224, *p* = 0.01, *η_p_*^2^ = 0.29). Follow-up *t*-tests indicated that VDT2 yielded steeper slopes compared to both the VDT0 (*p* = 0.04, Cohen’s *d* = 0.40) and VDT1 (*p* = 0.03, Cohen’s *d* = 0.24) conditions, indicating better performance in the condition, in which the last tactile stimulus was aligned with the disappearance of the visual stimulus (VDT2). No significant difference was noted between VDT0 and VDT1 (*p* = 0.73, Cohen’s *d* = 0.18). The comparison of two baseline conditions, i.e., VCAC in visual-auditory session and VCT0 in the visual-tactile session, yielded no significant difference (*F* (1, 27) = 0.08, *p* = 0.77, *η_p_*^2^ = 0.003).

## Discussion

The present study examined whether a temporal pattern imposed by sensory stimulation alters temporal prediction performance. In the visual-auditory session, visual and auditory oscillatory stimuli at a frequency in the delta-band significantly improved discrimination sensitivity relative to baseline, while decaying auditory stimuli significantly impaired performance. These effects were observed independently for both modalities, with no significant interaction. This suggested that a delta-band temporal pattern may enhance temporal prediction in this task both within the visual modality and across modalities via auditory stimulation. In the visual-tactile session, oscillatory visual stimuli improved sensitivity relative to decaying visual stimuli, supporting the interpretation that visual oscillatory stimulation facilitates temporal prediction, consistent with the visual-auditory conditions. However, rhythmic tactile stimuli failed to produce a similar behavioral benefit, suggesting that repeated sensory events alone are insufficient. Notably, the phase alignment of tactile stimuli relative to the visual stimulus critically determined performance: stimuli aligned to the visual offset (VDT2) yielded significantly better performance than those aligned to onset (VDT1) or absent completely (VDT0), providing behavioral support for the view that phase information contributes to temporal prediction.

The present findings suggest that oscillatory temporal patterns in the delta-band, induced by the sensory input, facilitate temporal prediction. The improvements relative to the condition in which stimulus strength was constant indicate that the critical factor was not simply the presence of additional stimulation, but the temporal organization of that stimulation. This was further supported by the absence of a significant difference between the unimodal (VCT0) and bimodal (VCAC) baselines. Notably, the improvement was not only attributable to the task-relevant visual modality but also to the auditory input, which suggests that such effects can operate across modalities, provided that the temporal pattern of sensory signals aligns with the underlying prediction process. At the same time, the finding that rhythmic tactile stimulation did not yield comparable benefits in the condition in which tactile stimuli were not aligned to the visual disappearance (VDT1), argues against the assumption that the behavioral benefits observed in oscillatory conditions were merely attributable to the presence of explicit temporal markers. Otherwise, providing repeated temporal events with the same frequency should have improved performance compared to the unimodal visual conditions, which was not the case. This indicates that a specific temporal profile of stimulation may be required to produce the behavioral enhancement. Alternatively, entrainment efficacy may be modality-dependent, potentially reflecting differences in the temporal resolution, precision in sensitivity, or cortical projection of these sensory systems. Taken together, our results are consistent with the hypothesis that temporal prediction benefits from an externally imposed oscillatory rhythm.

Furthermore, in the decaying condition, in which the stimulus disappearance was poorly defined, our findings indicated decreased discrimination sensitivity, in both the visual and auditory modalities. According to previous results^9^, the stimulus offset resets the phase of intrinsic delta oscillations, which then presumably serves as a reference point for temporal predictions. The present results support the critical role of phase information in the underlying delta oscillation for temporal predictions. The tactile findings are particularly informative in this regard: only the tactile stimuli aligned to visual offset (VDT2) improved performance relative to VDT1, in which tactile stimulation was aligned to stimulus onset. These results demonstrate that it is not merely the occurrence of rhythmic events that is relevant for successful temporal prediction, but the phase-specific temporal relationship between sensory input and the internal temporal profile.

Transcranial alternating current stimulation (tACS) has been widely used to entrain endogenous oscillations through externally applied electric fields^10–13,17^. The present study pursued an alternative approach, seeking to achieve comparable behavioral effects through sensory synchronization embedded within the stimuli themselves. Behaviorally, this approach was effective. The oscillatory stimulation in either the visual or auditory modality enhanced performance in a visual temporal prediction task, consistent with the effects reported for tACS-based entrainment^11,12^. Notably, cross-modal auditory entrainment produced performance comparable to, or even better than, that of unimodal visual entrainment, suggesting that the underlying timing mechanisms are not strictly modality-specific. This cross-modal efficacy offers a practical advantage: introducing oscillatory modulation in a modality separate from the task avoids potential interference with task-relevant stimulus features. Future neurophysiological studies using EEG or MEG could clarify the neural mechanisms underlying sensory entrainment and enable direct comparison with tACS-based approaches.

Several limitations of the study should be acknowledged. First, the present study is based on behavioral data only and therefore cannot directly demonstrate that the observed performance enhancements were mediated by entrainment of endogenous delta oscillations. The behavioral pattern is consistent with such an interpretation, but neurophysiological data are needed for definitive conclusion. Second, the absence of a general benefit of tactile stimulation in the delta frequency range indicates that the efficacy of external stimuli may depend on stimulus modality or on the detailed temporal profile of the stimulation. Neurophysiological studies are also needed to disentangle these possibilities. Third, we employed a fixed value in the delta frequency range in our experiment, drawing upon a previous study that utilized a comparable paradigm^9^. It is plausible that individual participants might display minor variations in their optimal frequencies within the delta range. Additionally, temporal prediction tasks using moving stimuli with different speeds might be reliant on oscillatory activity at other frequencies. Thus, it remains to be tested whether the precise frequency of the entrainment stimuli is relevant. Fourth, we found an asymmetry in the present results regarding sensory modalities: in the visual-auditory session, visually decaying stimulation did not significantly reduce performance relative to baseline, whereas its auditory counterpart did. A possible explanation is that the effect of visual decay was partially compensated by concurrent auditory oscillatory input. This interpretation is broadly consistent with results from the visual-tactile session, in which no auditory input was present and decaying visual conditions were associated with poorer performance. A 3 x 3 design with additional level of “constant”, besides “oscillating” and “decaying” in the current 2 x 2 design, would have allowed a cleaner dissociation of this effect. However, such an expansion would substantially increase the number of conditions and, consequently, the duration and demands of the experiment.

In summary, sensory oscillatory stimulation in delta frequency improved performance in several temporal prediction tasks; whereas decaying sensory stimulation impaired it. Depending on the specific phase of tactile stimulation, this impaired performance could be compensated. These results support the hypothesis that temporal prediction can be facilitated by an externally imposed oscillatory stimulation, either within or across sensory modalities. The importance of phase information in delta-band timing processes involved in predictive behavior was highlighted. More broadly, our findings suggest that structured sensory stimulation could be a useful tool for investigating and modulating temporal predictions across modalities.

## Methods

### Participants

A total of 32 healthy participants (21 female, 11 male) aged between 20 and 32 years (mean = 25.0 years, standard deviation = 3.2 years) were recruited for this study. All participants had normal or corrected-to-normal vision and were determined to be right-handed, as assessed by the Edinburgh Handedness Inventory^18^. Prior to participating in the experiment, all participants provided written informed consent to participate in the experiment in accordance with the ethical principles outlined in the Declaration of Helsinki. The study was approved by the ethics committee of the Medical Association Hamburg (PV7102).

All participants were invited to participate in two experimental sessions, one visual-auditory and one visual-tactile. The order of the two sessions was counterbalanced across participants. A total of 31 participants completed the visual-auditory session, and 29 participants completed the visual-tactile session, resulting in 28 participants completing both sessions.

### Stimuli and Procedure

#### Temporal prediction task

In this experiment, we tested whether sensory stimulation with different temporal patterns improve or impair behavioral performance in a temporal prediction task. For the purpose of this experiment, we used a temporal prediction task adapted from Daume et al. ^9^. In the task, a visual stimulus (i.e., an elongated oval) moved across the screen, disappeared behind an occluder and reappeared again. When the oval reappeared, participants were asked to judge whether it reappeared “too early” or “too late” in relation to their expectation.

#### Trial Structure

Each trial (Figure 1a) started with the presentation of the occluder, i.e., a rectangular area of greyscale white noise in the center of the screen, which was generated randomly and smoothed with a Gaussian filter. The occluder measured 5 degrees by 8 degrees visual angle (height by width) and had a red dot with a diameter of 0.12 degrees superimposed in its center, which served as the fixation point. The remaining area of the screen was a uniform grey background, which had the same brightness as the average of the occluder. Following a randomized inter-trial interval between 1.5 and 2.5 seconds, an oval (3 degrees by 0.8 degrees) appeared on either the left or right side of the screen (counterbalanced across participants) and moved towards the fixation point at a constant speed of 3.6 degrees per second. After approximately 2.9 seconds, the oval disappeared behind the occluder and reappeared after varying delays. The reappearance delays ranged from -0.933 to 0.933 seconds, in steps of 0.05 seconds, with negative values indicating earlier, and positive values indicated later, as compared to the exact “on time” reappearance. Note, the reappearance was never exactly on time. After reappearance, the oval continued moving for an additional 0.1 seconds. Next, the fixation point turned green, signaling participants to respond. Participants had to judge whether the reappearance of the oval was “too early” or “too late” relative to their expectation by pressing one of two buttons on a response box. Key mapping of responses was counterbalanced across participants. The next trial began either after the button press or 10 seconds of inactivity.

#### Sessions and Conditions

The task was performed in two sessions (Figure 1b): one Visual-Auditory (VA) session and one Visual-Tactile (VT) session. In the VA session, in addition to visual stimuli described above, auditory stimuli (A), i.e., pink noises, were presented, whereas in the VT session additional tactile (T), i.e., Braille vibration stimuli, were presented.

In the VA session, the stimuli oscillated (VO, AO), decayed (VD, AD) or remained constant (VC, AC), which resulted in five different conditions (VOAO, VDAO, VOAD, VDAD, VCAC). In the VCAC condition, the visual luminance and the auditory volume were always at their maximum values. In conditions with oscillating stimuli, the visual luminance and/or auditory volume were modulated sinusoidally, with the maximum luminance/volume equal to that in the VCAC condition and the minimum luminance/volume equal to the background. Participants experienced this as a flashing of light/sounds. The cycle period of the oscillation was 1.875 Hz, and the phase started at maximum and ended at a minimum. This resulted in 5.5 cycles from stimulus onset to its disappearance. In conditions with decaying stimuli, the maximum and minimum luminance/volume were set exactly the same as in the oscillating conditions, but decreased following a cubic function (i.e., faded away until no longer visible/audible).

Similarly, the VT session comprised six conditions (VCT0, VOT0, VDT0, VOT1, VDT1, VDT2). The baseline condition VCT0 and the oscillatory and decaying conditions (VOT0, VDT0) were unimodal visual. For all other conditions in this session, the visual stimuli were accompanied by tactile stimuli. For VOT1, VDT1, the tactile stimuli were aligned to the onset of the visual stimuli, whereas for VDT2, the tactile stimuli were aligned to the offset (i.e., disappearance) of the visual stimuli. This corresponded to the luminance peak and trough of the visually oscillating conditions respectively.

Prior to the task, participants received 120 trials of training (VCAC for the VA session and VCT0 for the VT session). Subsequently, all experimental conditions were presented in blocks of 120 trials (repeating all reappearance delays five times) in a random order. The duration of each block varied between 10-20 minutes, depending on the response speed of the participants. Participants were allowed to take a break and decide for themselves when to start the next block.

#### Hardware setup

The study was conducted in a dimly-lit, sound-attenuated cabin where participants were seated comfortably. Visual stimuli were presented on a BENQ LCD monitor located 60 centimeters in front of the participants, with a resolution set at 1920 x 1080 and a refresh rate of 60 Hz. Auditory stimuli were presented to participants through a 3M E-A-R TONE air-tube system, which were connected to the computer sound card. The maximum volume was adjusted based on the participant’s comfort and kept constant across all blocks. To present tactile stimuli, a Metec Braille piezo stimulator (Metec, Germany) with eight pins was attached to the index finger of the participant’s left hand. The eight pins were arranged in two columns with four rows, each 1 mm in diameter with a 2.5 mm spacing. For the purpose of the experiment, all eight pins moved synchronously together to provide a single tactile stimulus. The Braille stimulator was connected to a Texas Instruments board, which was controlled by output pulses from the computer’s parallel port. The on-phase of the tactile stimuli was constructed using 200 Hz vibration of the Braille pins, with the pins in the up-state 80% of the time in each cycle. These parameters were obtained in pilot tests by the experimenters to achieve optimal stimulus strength, which was kept consistent for all participants throughout the experiments. In the visual-tactile session, participants wore earplugs to prevent them from hearing the movement of the Brille pins. Participants responded to stimuli by pressing buttons on a button box connected to the stimulus PC. All stimuli were generated and delivered using the MATLAB-based software package Psychtoolbox^19^, which also collected and recorded responses. The software ran on a DELL personal computer equipped with Microsoft Windows 10.

#### Data acquisition and analysis

The results (Figure 2a) were plotted against physical delays, and more efficient detection was indicated by steeper rising slopes, meaning that smaller physical delay increases were required to achieve a given difference in the ratio of “too late” responses. The data were fit using the sigmoid function:

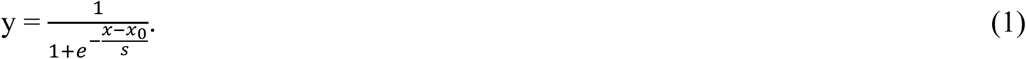

where *y* represents the percentage of “too late” responses; *x* represents the physical delays of reappearance to disappearance; *x_0_* represents the estimated “on time” delay, assuming a constant movement speed; and *s* represents the slope, which was submitted for further analysis (see below). A smaller value of *s* indicated a steeper slope, or better performance (Figure 2a).

The collected data were analyzed using MATLAB version 2020b, and subsequently processed with IBM SPSS software (version 26) for statistical analysis. Participants whose correct response rate to the (very easy) stimuli with a 2-second delay fell below 60%, were excluded from further analysis. Button presses within each block (condition) were converted to a ratio of “too late” responses for each delay, using the formula P (too late) = Number (too late) / (Number (too late) + Number (too early)). Any missing responses or button presses that did not correspond to either of the two buttons were omitted from analysis. The data were then fit for each block (condition) using equation (1). The time delays to physical on-time reappearance, were taken as dependent variable.

The fitted parameter *s* was used to indicate the slope of the curve and represent the sensitivity of discrimination. Any data associated with a goodness-of-fit (adjusted *r*-squared) value less than 0.5 were excluded. Participants whose data exceeded two times of standard deviation from the mean, were also excluded. In the end, 26 participants in the VA session and 27 participants in the VT session, were formally analyzed.

#### Statistical Analysis

For the VA session, the fitted parameter values of *s* from VDAD, VOAD, VDAO, and VOAO were normalized by dividing *s* of each condition by *s* of the constant condition (VCAC). We performed a 2 x 2 ANOVA with two within-subject factors: visual and auditory, each consisting of two levels (oscillating vs. decaying). When the assumption of sphericity was violated, Greenhouse-Geisser correction was applied. In multiple comparison cases, Šidák-correction was employed. The averages of all involved conditions of a certain level of a factor, e.g., the average of VDAD and VOAD serve as auditory decaying (AD), were used for their comparisons to baseline with *t*-tests. Effect size was reported alongside the *p*-values, expressed as partial eta-squared (*η_p_*^2^) or Cohen’s d (*d*), depending on the type of statistical test.

Similarly to the VA session, in the VT session parameter *s* was normalized to the unimodal condition VCT0. We performed a baseline comparison similar to the VA session using *t*-tests. To analyze the two factors visual stimulation (oscillating vs. decaying) and tactile stimulation (on vs. off), we performed a 2 x 2 ANOVA. To analyze the effects of phase alignment of the tactile stimuli, we performed a 1 x 3 repeated measures ANOVA with one within-subject factor of tactile stimuli type: tactile stimulus off (VDT0), in-phase to vision (VDT1), and anti-phase to vision (VDT2). The parameter *s* of two baseline conditions was also compared with pair-wise *t*-tests, to investigate the effect of the additional auditory stimuli without oscillation.

## Acknowledgements

A.K.E. acknowledges support for this work from the DFG (TRR169-261402652-B1/B4/Z2). We thank Karin Reimann for participant recruitment and Rebecca Burke for helpful discussions of the results. The information and views expressed in this paper are purely those of the authors and do not necessarily reflect the official opinion of the European Commission or the European Research Council Executive Agency.

